# Thousands of human non-AUG extended proteoforms lack evidence of evolutionary selection among mammals

**DOI:** 10.1101/2022.05.02.490320

**Authors:** Alla D. Fedorova, Stephen J. Kiniry, Dmitry E. Andreev, Jonathan M. Mudge, Pavel V. Baranov

**Author notes:** corresponding authors Alla D. Fedorova.

## Abstract

The synthesis of most proteins begins at AUG codons, yet a small number of non-AUG initiated proteoforms are also known. Here we used publicly available ribo-seq data with phylogenetic approaches to identify novel, previously uncharacterised non-AUG proteoforms. Unexpectedly we found that the number of non-AUG proteoforms identified with ribosome profiling data greatly exceeds those with strong phylogenetic support. We identified an association between proteoforms with alternative N-termini and multiple compartmentalisation of corresponding gene products. In dozens of genes N-terminal extensions encode localisation signals, including mitochondrial presequence and signal peptides. While the majority of non-AUG initiated proteoforms occur in addition to AUG initiated proteoforms, in few cases non-AUG appears to be the only start. This suggests that alternative compartmentalisation is not the only function of non-AUG initiation. Taking a conservative approach, we updated annotation of several genes in the latest GENCODE version in human and mouse where non-AUG initiated proteofoms are supported by both, ribosome profiling and phylogenetic evidence. Yet, the number of such extensions is likely much higher. Thousands of non-AUG proteoforms supported only by ribosome profiling suggest that they may evolve neutrally. Indeed, expression of some may not be consequential, i.e. when N-termini is processed or they have identical biochemical properties. Nonetheless they may contribute to immune response as antigen sources. It is also possible that some proteoforms accrued useful functions only recently and evolved under purifying selection in a narrow phylogenetic group. Thus, further characterisation is important for understanding their phenotypical and clinical significance.

## Introduction

The current paradigm of translation initiation in eukaryotes follows the scanning mechanism wherein the preinitiation complex (PIC), assembled on the small ribosomal subunit (40S) and containing initiator Met-tRNAi (methionyl tRNAi), scans the mRNA 5’ leader for an AUG codon in a suitable context using complementarity to the anticodon of Met-tRNAi. The first AUG codon entered the peptidyl-tRNA (P) site of the 40S subunit is usually employed as the start codon, but it can be missed in unfavourable surrounding context. When optimised, this ‘Kozak’ context has a purine at position -3 and guanine at +4 position relative to the AUG (+1 position). When the first AUG codon is skipped due to weak Kozak context, the next AUG codon can be used. This phenomenon is known as leaky scanning (Hinnebusch 2014; Kozak 1980).

Although it was long believed that the synthesis of eukaryotic proteins initiates at an AUG start codon, translation initiation at codons differing by 1nt from AUG (near cognate) have been documented in early 80s, albeit with much lower efficiency (Anderson and Buzash-Pollert 1985; Hann et al. 1988; Peabody 1989; Kozak 1989). It occurs in spite of the near cognate codon mispairing with the anticodon of Met-tRNA. This mispairing can be tolerated only during initiation because it is the only stage when an incoming Met-tRNAi is bound directly in the ribosomal P-site (Simonetti et al. 2008; Ramakrishnan 2002). Unlike the A-site, where mRNA:tRNA interactions are thoroughly monitored by the decoding centre (Ogle et al. 2001), the P-site is more promiscuous to mismatches in the codon:anticodon duplex (Potapov et al. 1995; Baranov et al. 2004; Svidritskiy and Korostelev 2015; Gallant and Masucci, 2000). Near cognate triplets such as CUG, GUG, UUG, AUA, AUU, AUC, ACG have been shown to be recognised as starts at frequencies of ∼1-10% of AUG in the optimal context depending on the gene and study, while AAG and AGG are essentially not recognised (Peabody 1989; Kearse and Wilusz 2017). However, there is an astonishing example of a highly efficient CUG initiation conserved in mammals. The CUG codon is located in the 5’ leader of *POLG* gene which encodes the catalytic subunit of the mitochondrial DNA polymerase. The efficiency of initiation at this CUG is comparable (∼60-70%) to that at an AUG in the optimal context (Loughran et al. 2020). It results in translation of a 260-triplet-long overlapping open reading frame (Khan et al. 2020) called POLGARF, its functional role is suggested to be involved in the extracellular signalling (Loughran et al. 2020). Another example of a very conserved near-cognate initiation is the *EIF4G2* gene, whose translation is initiated at a GUG. It encodes a paralog of eIF4F complex subunit eIF4G, but lacks the binding site for the cap-binding subunit eIF4E. The GUG-initiation rate for *EIF4G2* is unusually high, ∼30% compared to an AUG-mutant version of the same *EIF4G2* expression plasmid (Imataka et al. 1997; Tang et al. 2017). Initiation efficiency from a non-AUG start may be enhanced by secondary structure elements starting approximately 15 nt downstream of the non-AUG codon (Kozak 1990). The Kozak context for non-AUG codons is similar to AUG and also important for initiation efficiency (Kearse et al. 2016).

Due to leaky scanning, translation initiation from non-cognate start codons may result in extended proteoforms in addition to a proteoform resulting from initiation at the “main” downstream AUG (Fig.1A). Genes with multiple non-AUG initiated proteoforms are known, e.g. human tumour suppressor PTEN with firstly identified CUG-initiated N-terminally extended proteform (Ivanov et al. 2011; Hopkins et al. 2013), then more abundant AUU-initiated proteoform and two additional CUG-initiated proteoforms (Tzani et al. 2016). All of them retain the ability to downregulate the PI3K pathway (Liang et al. 2017). Another possibility is translation initiation from non-cognate start codon downstream of AUG which leads to a truncated proteoform e.g. in *MRPL18 (Zhang et al. 2015)* and *ASCT2 (Tailor et al. 2001)*. Cases of exclusive translation initiation (Fig.1B) at non-AUG codons have also been reported, e.g. already mentioned mammalian *EIF4G2* (Takahashi et al. 2005), human *TRPV6 gene* (and its mouse *Trpv6* homolog) generates a single ACG-initiated TRPV6 protein, human STIM2 initiates exclusively at an UUG start codon (Williams et al. 2001), human *TEAD1* exemplifies translation initiation at AUA start codon (Xiao et al. 1991). In total, more than 60 instances of non-AUG-initiated proteoforms have been reported (Ivanov et al. 2011; Van Damme et al. 2014).

**Fig. 1.**
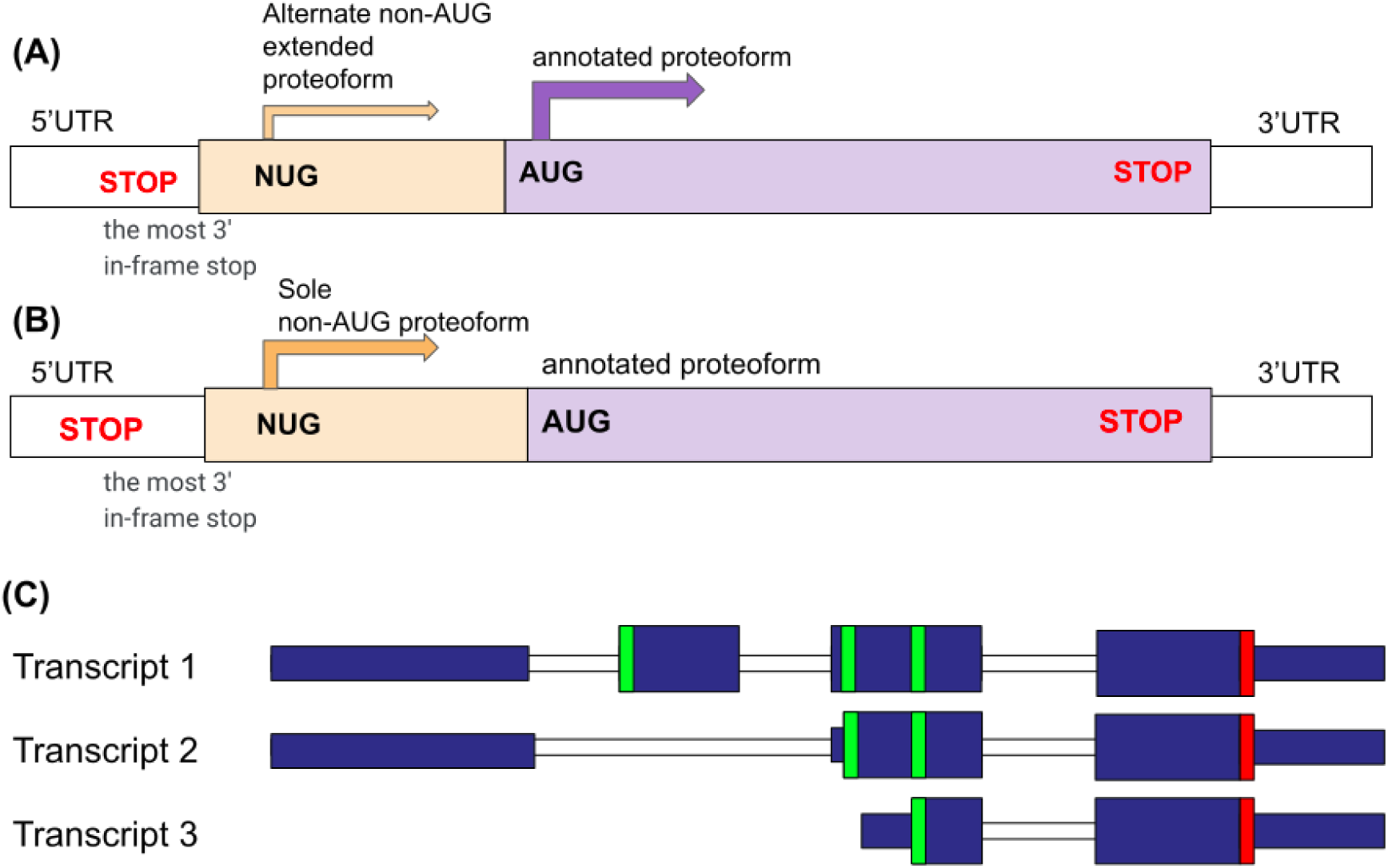
Transcript structure for genes encoding non-AUG initiated proteoforms **(A, B)** and alternative splicing and transcription start sites as sources of alternative N-termini **(C)**. **(A)** Transcript with both major annotated proteoform and N-terminally extended non-AUG initiated proteoform. **(B)** Transcript with sole non-AUG initiated proteoform. **(C)** Transcript 1 has an extra CDS exon relative to transcript 2, while transcript 3 has a different transcription starts. As a result, non-coding parts of first CDS exons in transcripts 2 and 3 exhibit protein coding evolution.

Detection and annotation of non-AUG initiated proteoforms is clinically important since their misregulation may lead to multiple human diseases including neurodegenerative disorders and cancer progression. For instance, *FGF2* controls cell proliferation, differentiation and angiogenesis and it has a canonical AUG-initiated proteoform which is mostly cytoplasmic or secreted. At least four upstream CUG-codons can be used to generate longer isoforms that localise to the nucleus therefore causing cell immortalisation (Arnaud et al. 1999; Bugler et al. 1991). The *MYC* proto-oncogene regulates cell proliferation and transformation. Two proteoforms are generated from *MYC* using a canonical AUG codon and an upstream in-frame CUG codon. The CUG-encoded proteoform becomes more prevalent during the limited availability of amino acids when the density of cells increases and its overexpression was shown to inhibit the growth of cultured cells. Inactivation of this proteoform is observed in Burkitt’s lymphomas suggesting that the inability to generate this CUG-initiated proteoform may provide a selective growth advantage (Hann and Eisenman 1984; Hann et al. 1988).

Detection of extended non-AUG proteoforms can be carried out by various approaches. Bioinformatics methods include comparative genomic analysis where multiple nucleotide sequence alignments analysed for the presence of substitution patterns typical for protein coding evolution e.g. reduced rate of non-synonymous substitutions relative to non-synonymous measured as their ratio, d_N_/d_S_ or Ka/Ks (Ivanov et al. 2011). Western blots, ribosome profiling and proteomics can serve as experimental support of predicted extensions. The identification of non-AUG proteoforms can be assisted with application of machine learning methods (Reuter et al. 2016).

## Results

### Detecting purifying selection upstream of annotated coding regions

It is well established that purifying selection is a typical evolutionary signature of protein-coding sequences. One would expect that if translation initiates upstream of the annotated start codon, this upstream region should evolve as a protein-coding sequence. We used this indicator in order to identify N-terminally extended proteoforms in the human genome by means of PhyloCSF score. PhyloCSF is a method for assessing the evolutionary protein-coding potential of a genomic region based on multiple sequence alignment (Lin et al. 2011). However, the protein-coding evolution of an upstream region does not necessarily mean that the underlying mechanism is the non-AUG initiation or alternative transcription (see Fig.1C).

Yet another possible mechanism is Stop Codon Readthrough (Baranov et al. 2015). When a uORF is in the same reading frame with the CDS and there are no stop codons between the uORF stop codon and the annotated CDS start, ribosomes reading through the uORF stop codon could produce a protein encoded by the fusion of the uORF and the extended CDS ORF. Quite noticeable (∼17%) level of readthrough produces C-terminally extended form of human opiate receptor OPRL1 (Loughran et al. 2017), while examples where readthrough mediates N-terminal extensions have not yet been reported.

Such an extension could also be generated with Programmed Ribosomal Frameshifting (Atkins et al. 2016). In antizyme genes, the CDS is organised in two overlapping ORFs, where ribosomes initiating at the first ORF shift reading frame at its stop and enter +1 ORF that encodes most of the protein. In the absence of the prior knowledge only the second antizyme ORF would be encoded as protein coding with an internal AUG annotated as a start. This is exactly what happened during the annotation of *S. cerevisiae* genome (Palanimurugan et al. 2004). Similarly, in a recent study (Ivanov et al. 2020) previously annotated yeast ORF YPL034w (ORF2) has been shown to encode only a part of the protein that is synthesised as a result of initiation at uORF followed by +1 frameshifting.

Determining the exact start of non-AUG extensions in practice can be quite challenging because there might be several non-AUG starts upstream of AUG like in the case of PTEN. So we focused on predicting the genes which are most likely to have alternative extended non-AUG proteoforms irrespective of our ability to identify specific locations of start codons. The final set of candidates (PhyloSET, see Supplemental Table 1) was obtained via following steps (see Methods for further detail) shown on Fig.2, steps 1-4a. PhyloCSF score is ranging from 0.1452 (*NRXN1*) to 2693.8893 (*CCDC8*) with median value 155.9191 (*TRPC1*). PhyloCSF scores per codon were calculated to observe how the selection changes over the selected upstream region. Ideally, we would expect that the start of extension is clearly separated from the non-coding sequence, in other words, PhyloCSF score becomes positive at the border between extension and preceding non-extension part as it happens for *CCDC8* gene (Fig.3A). However, for the majority of genes no such clear change in scores can be spotted perhaps because evolutionary selection on N-terminus is relaxed in comparison with internal parts.

**Fig. 2.**
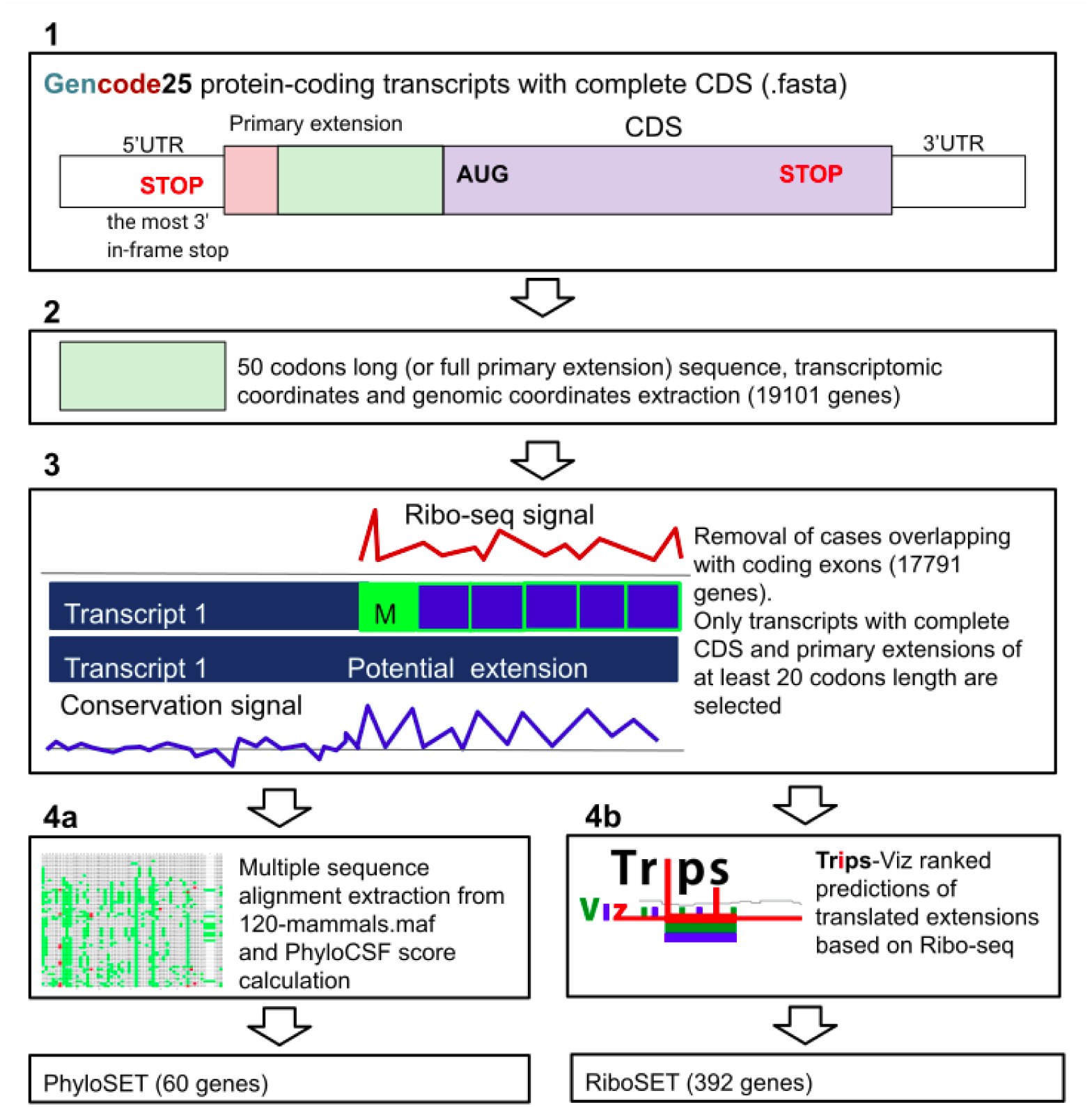
Scheme of the pipeline. (**Step 1**) GENCODE v25 protein-coding transcripts with complete CDS were used to extract primary extension; (**step 2**) 50-codons-long extension (or the whole primary extension if it is shorter than 50 codons) was obtained; (**step 3**) transcripts for which extensions overlap with coding exons in any reading frame were excluded; (**step 4a**) PhyloCSF score was calculated for 50-codons-long set of extensions using 120-mammals multiple sequence alignment; strictly positive threshold led to a set of 60 genes which we called PhyloSET; (**step 4b**) the retrieval of the ranked extensions predicted based on ribosome profiling data in Trips-Viz browser that composed RiboSET.

**Fig. 3.**
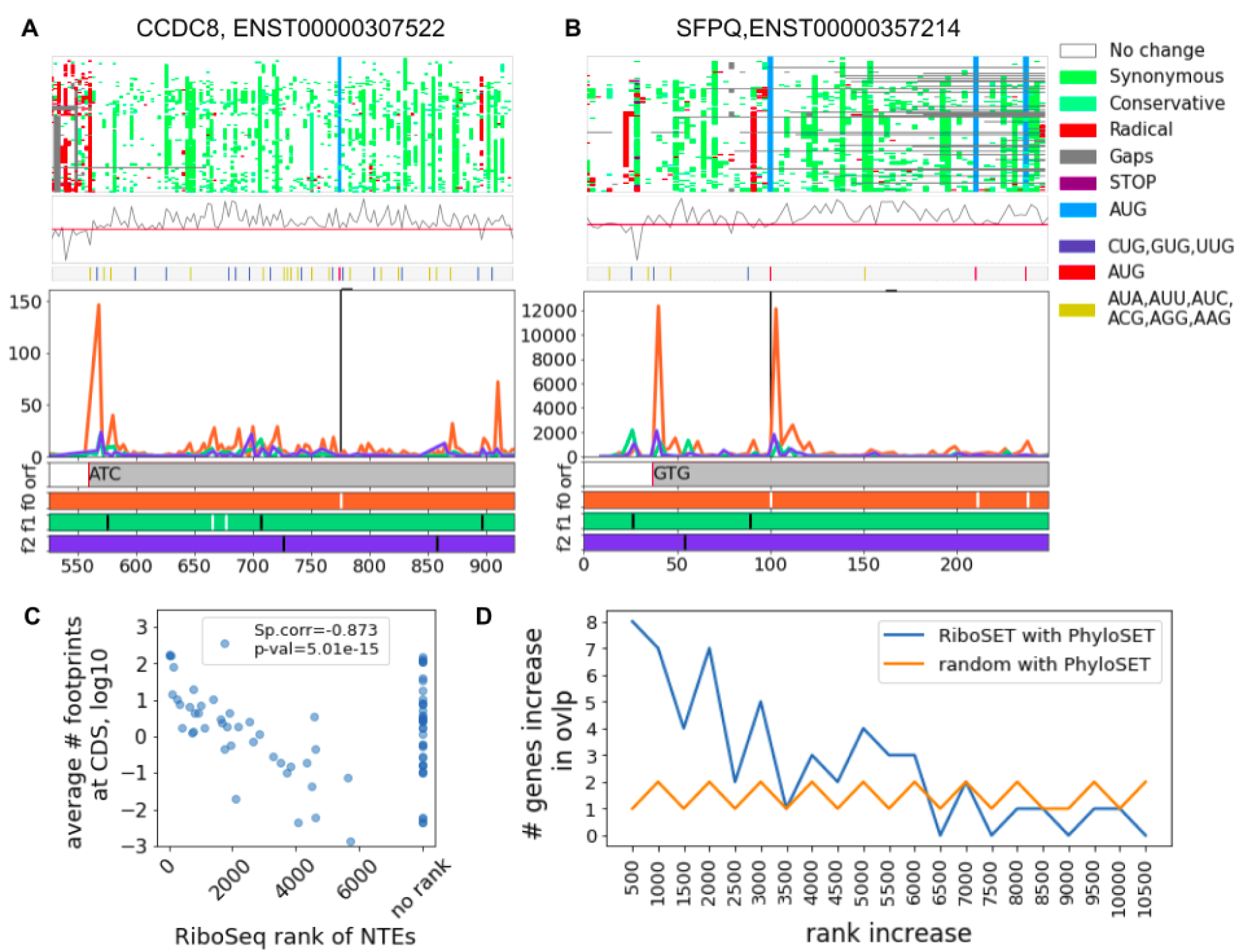
Genes found in both PhyloSET and RiboSET (**A** - *CCDC8*, **B** - *SFPQ*), genomic intervals including primary extension and the first 50 codons of CDS are shown. Each subplot contains three parts. Top plot is a colour coded codon alignment (each tile represents a codon, green-highlighted codons represent synonymous and conservative substitutions, red ones - radical substitutions, grey one - indels, purple - stops, blue - AUGs; adapted from CodAlignViewю Middle plot represents the PhyloCSF score per codon (the bottom colour bar shows positions of potential start codons: red shows AUG, blue represents XUG and yellow is used for remaining near-cognate start codons). Bottom plots show Trips-Viz subcodon Ribo-seq profiles with the densities of ribosome footprints differentially coloured based on the supported reading frame. The colours are matched to the reading frames in ORF plot at the bottom where AUG codons are depicted as white and stop codons as black dashes. Black vertical lines indicate the starts of the annotated CDS. Grey bars correspond to extended CDS initiated at the proposed non-AUG starts. **C**. Average footprint density at CDS of PhyloSET genes (in log10 scale) compared to the Ribo-Seq rank of N-extensions in them; ranks are not assigned below rank 10470. Spearman correlation is calculated for genes with known rank (corr=-0.873, p-value=5.01e-15). **D**. The size of the increase in the number of genes between PhyloSET and RiboSET overlap depending on RiboSeq NTE threshold. The highest rank assigned with Trips-viz is 10470. The distributions (in the range from 500 to 6500 rank, where they overlap) are compared with Mann–Whitney U test (p-value=0.0016, statistic=28.5).

### Detecting translation upstream of annotated coding regions with Ribo-seq and proteomics data

Another set of candidates (RiboSET) was selected solely based on ranked translated extensions predicted using ribosome profiling data with Trips-Viz, (Fig.2, steps 1, 2, 3, 4b, Supplemental table 2) (Kiniry et al. 2019, 2021). Ribosome profiling is a method based on deep sequencing of ribosome-protected mRNA fragments which was introduced by Nicholas Ingolia and Jonathan Weissman (Ingolia et al. 2012). This set was generated with a ranking procedure for detection of translated regions implemented in Trips-Viz browser which is described in Methods. In the absence of objective threshold for discriminating genuine translation from biological and technical noise, we decided to incorporate 500 top scoring extensions into RiboSET which upon further filtering (Methods, Fig.2) was reduced to 392 genes. Although important to mention that the actual number of translated non-AUG extensions is much higher (some extensions ranked below 5000 are reliably translated). Proteomics data available in Trips-Viz (Kiniry et al. 2021) supported extensions in 90 genes in this RiboSET (18 in PhyloSET). Only 8 genes (*CCDC8*, *CYTH2*, *FXR2*, *H1FX*, *HNRNPA0*, *MARCKS*, *RPTOR*, *SFPQ*) are common between PhyloSET and RiboSET (Fig.3A, B, Fig.S1 in Supplemental text 1).

The small overlap between PhyloSET and RiboSET requires an explanation (more details are in discussion). Genes may occur in PhyloSET exclusively either because they are not expressed in the cells for which Ribo-seq data are available or because the 500 top ranking threshold is too conservative. To explore the first possibility we studied a relationship between rank of predicted extension and CDS coverage - average number of footprints at CDS. We observed that the lower the average number of footprints at CDS, the lower extension is ranked (Fig.3C). To show whether a threshold of 500 top ranking candidates is too conserved, we explored the size of the overlap increase (we use step = 500) between PhyloSET and RiboSET depending on ranking threshold, see Fig 3D. It appears that until the rank reaches 6500, there is a statistically significant (p = 0.0016, Mann–Whitney U test) increase in the overlap size in comparison with what would be expected by chance suggesting that Ribo-seq signal above our selected threshold has a clear positive value at predicting genuine N-terminal extensions. Genes with high CDS coverage but with no rank may still have extension translated under certain conditions (examples can be found in Fig.S2, Supplementary text 1).

### In-frame and out-of-frame AUGs in theoretical extensions of PhyloSET and RiboSET

Assuming that during genome annotation it is the first AUG codon that is usually annotated as a start (in the absence of additional experimental evidence overriding this rule) we were surprised to find in-frame AUG codons in the N-terminal extensions of twelve genes from RiboSET and PhyloSET. In 8 genes (*DUSP5*, *PDCD6*, *CCDC127*, *EID2*, *DHX33*, *SLC17A5*, *TMX3*, *MXD4)* we found that their 5’ leader are longer in GENCODE v25 in comparison to RefSeq ones which do not possess such upstream non-annotated AUGs. No evolutionary or translational signatures advocate for use of such in-frame AUGs in these genes. In case of *DUSP5* and *PDCD6*, upstream AUG is located very close to the 5’ end of the transcript (4nt and 5nt) and may not be recognised by the pre-initiation ribosomal complex. Transcripts in GENCODE v25 and v35 and RefSeq for *TK1* gene have an upstream AUG and a longer 5’ leader with quite decent conservation (high PhyloP score) although RNA-seq data (available in GWIPs-viz (Michel et al. 2014) does not seem to support such a long 5’ leader. Of note, high PhyloP score could be due to a conserved promoter that is not a part of transcript as was recently shown for *Hoxa* genes (Ivanov et al. 2022). According to Trips-viz ribo-seq profile, the upstream AUG does not seem to have any translation signal (Fig.S3, Supplemental text 1). Trips-viz Ribo-seq profiles of *C1GALT1, LY6K and DHFR* look like there might be an AUG-extension (Fig.S3, Supplemental text 1), although AUGs in *DHFR* are located in the overlap with coding exons of another gene (*MSH3*) on a different strand (Fig.S4, Supplemental text 1).

Trips-viz predicted 3 more genes with AUG extensions. *STIM2* (STromal Interaction Molecule 2) has clearly translated AUG-extension according to Trips-viz Ribo-seq profile while it has only UUG proteoforms in GENCODE v35 and the latest RefSeq in humans. In contrast, in mice both AUG- and UUG-proteoforms are annotated. *PTPRJ* (Protein Tyrosine Phosphatase, type J) has also been shown to possess AUG-extension according to Trips-Viz predictions. An upstream AUG initiation has been described in Karagyozov et al. 2020 (Karagyozov et al. 2020). *AP3S1* (Adaptor Related Protein Complex 3 Subunit Sigma 1) is the last candidate with AUG-extension predicted by Trips-viz (Fig.S5, Supplementary text 1).

We also addressed the occurrence of out-of-frame AUGs between non-AUGs and annotated AUGs. Out-of-frame AUG are a source of uORFs which may be involved in the regulation of relative proteoform synthesis. Translation of an out-of-frame uORF is likely to inhibit the synthesis of the extended proteoform (especially when it overlaps the extension), but may be less detrimental to the translation initiation at annotated AUG which ribosome could access via reinitiation (Gunišová et al. 2018; Skabkin et al. 2013). On the contrary, initiation at out-of-frame AUG located within the extension is unlikely to affect the synthesis of the longer proteoform but would inhibit the synthesis of the shorter one. uORFs is a rich source of versatile mechanisms for gene-specific regulation at the translation level in response to specific conditions (Andreev et al. 2015b, 2015a; Starck et al. 2016; Mueller and Hinnebusch 1986) and it is likely that they may also be used for regulation of a ratio between proteoforms in a similar manner. In RiboSET, 57 genes (57 transcripts) have at least one out-of-frame AUG codon in their theoretical extensions. In PhyloSET, 24 genes (36 transcripts) have at least one out-of-frame AUG codon in their theoretical extensions. We explored their Ribo-seq profiles in order to spot translated uORFs. In some genes, uORFs are located within translated extension (e.g. *FAM102B*, *PIEZO1, SNRNP25*, *FBXL3*), while there are also examples where uORF is situated upstream of translated extension (e.g. *CDR2*, *PPP1R14B*, Fig.S6, Supplemental text 1).

### Comparison with previously identified non-AUG proteoforms

We also wanted to know how well translation detected with ribosome profiling concords with phylogenetic approaches. We performed comparisons of a set of predicted proteoforms from the previous study (Ivanov et al. 2011) with PhyloSET and RiboSET. In brief, the study utilised alignments of human and mouse RefSeq transcript sequences which resulted in prediction of 59 genes with evolutionary conserved extensions. PhyloSET and RiboSET are meant to have only new non-AUG proteoforms which have not been described in GENCODE v35 and the latest RefSeq annotation (due to exclusion of overlapping coding exons). In GENCODE v35, 24 non-AUG proteoforms from study (Ivanov et al. 2011) have been annotated; 28 genes have not been annotated with non-AUG proteoforms and retained intact extension sequences detected in the previous study and 7 genes remained without annotated near-cognate initiated proteoforms or intact extensions (*HELZ2, ANKRD42, WDR26, ZFP62, C1QL1, PTEN, TIAL1*, where *WT1* was shown to be annotated and intact extension in *TIAL1* still corresponds to only nonsense-mediated decay transcript, (Supplementary tables 3A-C). Among 28 genes there were 4 genes found in PhyloSET and 2 genes in RiboSET. Among 7 genes 1 gene and 0 genes were shown in PhyloSET and RiboSET correspondingly (Fig.4, A).

**Fig 4.**
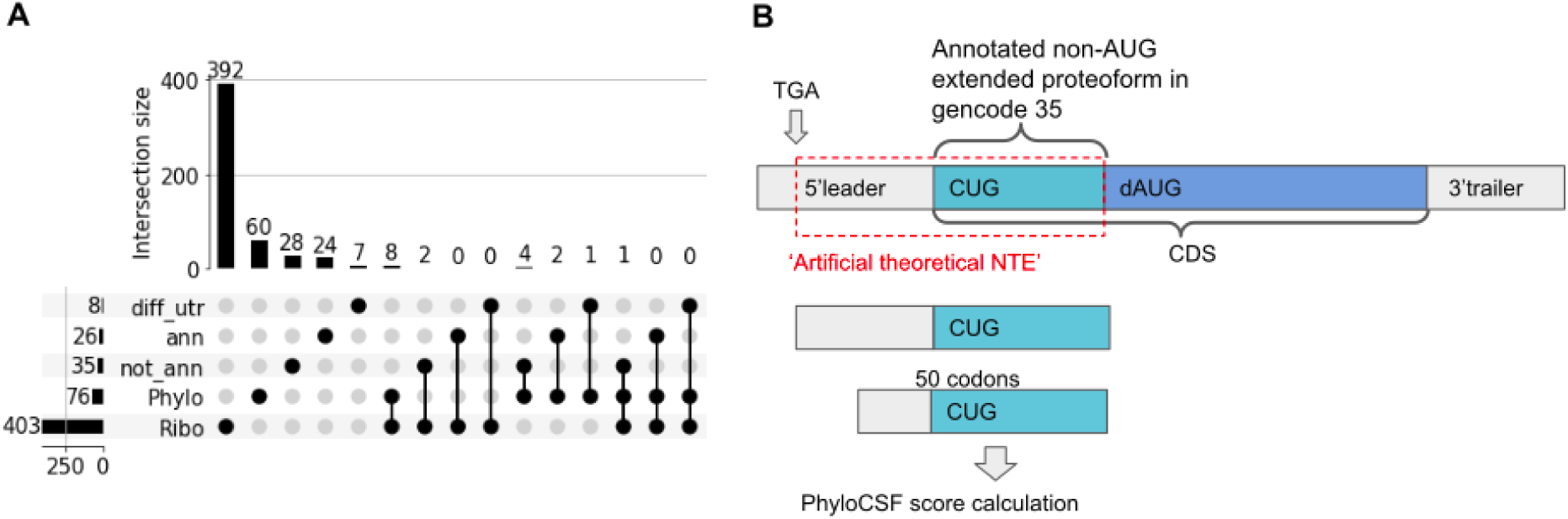
**A.** Comparison of RiboSET and PhyloSET with genes from Ivanov et al 2011 [21]. ‘Phylo’ - PhyloSET, ‘Ribo’ - RiboSET, ‘ann’ - genes with annotated non-AUG extensions in GENCODE v35, ‘not_ann’ - genes which non-AUG extensions are not annotated in GENCODE v35 (5’leader is the same as in previous study), diff_utr - genes which non-AUG extensions are not annotated in GENCODE v35 (5’leader is different). **B**. Re-identification of non-AUG N-terminal extensions in 24 genes from study Ivanov et al, 2011 which have been annotated in GENCODE v35. ‘Artificial theoretical NTE’ starts from the most 3’ in-frame stop codon and stretches till the first downstream ATG right after non-AUG. PhyloCSF score is calculated for the first upstream 50 codons of theoretical extension.

We also decided to test whether a PhyloCSF-based approach is able to re-identify those 24 genes which have been already annotated in GENCODE v35. First, we created a set of transcripts with non-AUG proteoforms for these genes where the start of CDS is moved to downstream AUG (Fig.5, B). It turned out that the PhyloCSF score is positive for less than half of genes (11/24, Supplemental table 3D). The remaining 13 genes have not shown positive PhyloCSF score. It might be explained by the extended part being much shorter than 50 codons which may have led to an excessive codons’ impact on negative scores (*MYC, YPEL1, HCK, TRPV6*).

**Fig. 5.**
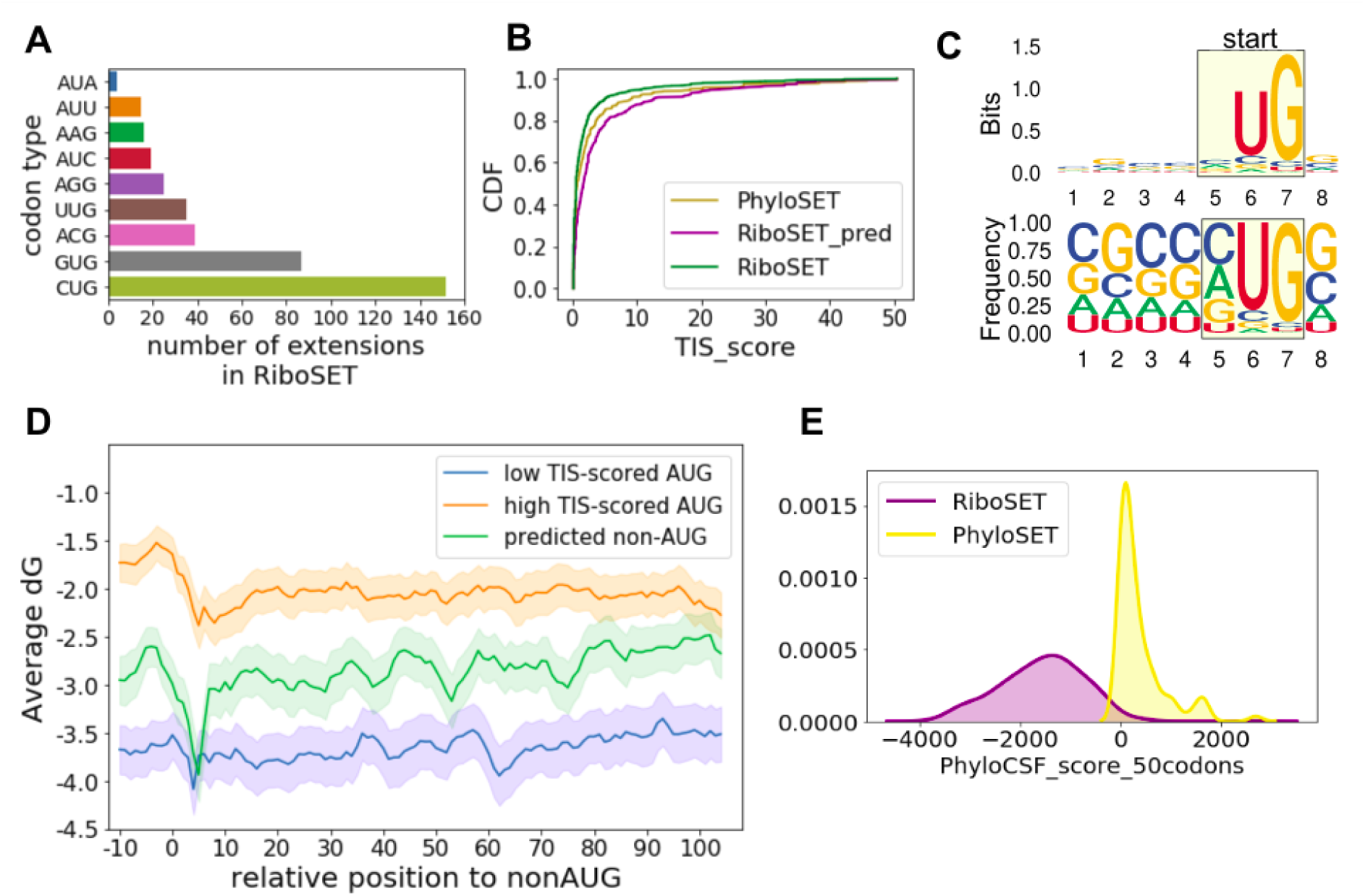
Characterisation of predicted non-AUG initiation codons. **A**. Distribution of start codon types predicted by Trips-viz (RiboSET). **B**. Cumulative distribution functions of TIS scores for all non-AUG codons in theoretical NTE, RiboSET (green), all non-AUG codons in theoretical NTE, PhyloSET (yellow), Trips-viz predicted non-AUG starts, RiboSET (purple). **C**. TIS sequence logo and frequency plot of non-AUG starts predicted by Trips-viz. **C**. TIS sequence logo and frequency plot of non-AUG starts predicted by Trips-viz. **D**. The stability (dG, Gibbs free energy) of mRNA secondary structure downstream of start codons within 22 nt window ‘0’ corresponds to the start codon. 392 starts from RiboSET (green line), sample of 400 AUG starts with high-scored TIS (blue line), sample of 400 AUG starts with low-scored TIS (orange line). Lines are mean values across genes with 95% confidence interval. **E**. Distribution of PhyloCSF score for RiboSET and PhyloSET genes.

We revisited Ribo-seq profiles for these 24 annotated in GENCODE v35 genes with non-AUG extensions (Fig. S7, S8, Supplementary text 1). For 7 genes there was lack of Ribo-seq data coverage for the entire mRNA (*FNDC5*, *NR1I2*, *PRPS1L1*, TRPV6, HCK, YPEL1, OAZ3). Extensions are clearly supported by Ribo-seq data in 16 genes including: *WT1*, *TEAD4*, *GTF3A*, *NPW*, *CACNG8*, *DDX17*, *VEGFA*, *R3HCC1*, *MYC, RNF187*, *NFKBID*, *YPEL2, BAG1* (transcript has no 5’leader), *EIF4G2* (although there is likely to be an additional non-AUG initiation upstream), *TEAD3* (there is also likely to be non-AUG upstream start site), *SP3* (another upstream site of non-AUG initiation can be seen). For *KCTD11*, although even if there is a Ribo-seq signal upstream of AUG, it is not clear whether extension starts where it is annotated.

In addition, we also compared our gene sets with Van Damme et al study (Van Damme et al. 2014) which identified 17 human genes with non-AUG N-terminal extension using ribosome profiling and N-terminal proteomics (Supplemental table 3E). Among 17 candidates, two genes from PhyloSET (*FXR2, HNRNPA0*) and seven genes from RiboSET (*NARS, HDGF, HNRNPA0, FXR2, SYAP1, KAT7, BAG6*) were present. We also compared RiboSET and PhyloSET gene lists with extensions detected with the N-terminal-peptide-enrichment method from the study Yeom L. et al (Yeom et al. 2017). Only 1 gene from PhyloSET and 17 genes from RiboSET were present among 171 genes with N-terminal extensions (Supplemental table 3F).

Of note, there was a significant discrepancy between RefSeq and GENCODE gene annotations in *PTEN*. In the latest RefSeq mRNA the CUG-initiated proteoform is annotated correctly, while in GENCODE v35 this proteoform has not been annotated yet and 5’ leader of the only one available transcript ENST00000371953. This can be explained by the incorrect sequence of the reference genome (assembly GRCh38) - it has the variant which is known as NC_000010.11:g.87864104delT and its global minor allele frequency in 1000 Genomes 0.00000 (T). This variant introduces frameshift into the 5’leader of a transcript thus disrupting the sequence of CUG-extension. In RefSeq gene annotation this variant is cut from transcript sequence and shown as 1nt-intron in Genome Browser (Fig. S9, Supplemental text 1).

### Characterisation of genes with predicted non-AUG initiation

Firstly, we studied the distribution of start codon type across starts predicted by Trips-viz in RiboSET. As expected, the most frequent non-AUG start in extensions was CUG. This was followed by GUG and ACG (Fig.5A). In RiboSET only one non-AUG initiation site per transcript is predicted based on internal probability ranking relying on features associated with the intensity of Ribo-seq signal. Therefore, one would expect that the initiation efficiency of such starts may be facilitated by certain features including the optimality of Kozak context and downstream secondary mRNA structures. It has been shown that certain non-AUG start codons with the appropriate sequence context can initiate translation comparable to that of AUG start codons (Diaz de Arce et al. 2018). The efficiencies of TIS (Translation Initiation Site) including start codon and four positions upstream and downstream were previously measured with FACS-seq. For this purpose, a library of fluorescent reporters under control of all possible contexts surrounding near-cognate initiation codons was transfected into cells and then cells were sorted based on fluorescence. The efficiency of a specific TIS was measured based on its enrichment within a specific fraction and scaled relative to optimal TIS (CACC<u>AUG</u>G) efficiency score set to 100 (Diaz de Arce et al. 2018) (the scores of non-AUG starts were found in the range from 0.2 to 50.4). We compared TIS scores of predicted non-AUG starts from RiboSET (Supplemental table S4) to all other non-AUG codons in theoretical extensions in RiboSET and PhyloSET (Fig.5B). It is clearly seen that predicted non-AUG starts in RiboSET have more favourable initiation contexts in comparison to all theoretical non-AUG codons in primary extension sequences thus endorsing Trips-viz-based prediction method (Kolmogorov-Smirnov 2-sample test, p-value 1.19e-18, statistic=0.245).

The next step was to assess the stability and presence of mRNA secondary structures located downstream of predicted start codons. It has been shown that a strong RNA secondary structure located downstream of the initiation site significantly increases the efficiency of initiation at non-AUG codons (Kozak 1990; Kochetov et al. 2007). We selected 400 genes with high-scored AUG-containing TISs and 400 genes with low-scored AUG-containing TISs as well as all predicted nonAUG TISs from RiboSET (392 genes). We then used RNAfold (Gruber et al. 2008) to calculate free energy of predicted RNA secondary) within a sliding window of 22nt with the step of 1 nt in the region surrounding potential start codon (10 nt upstream and 100 nt downstream), see Fig.6D. As expected, more stable mRNA secondary structures were present on transcripts with less optimal TIS codons. PhyloCSF score distribution for RiboSET genes is clearly skewed towards large negative values arguing for massive non-conserved non-AUG translation in 5’ leaders (Fig.5E).

**Fig. 6.**
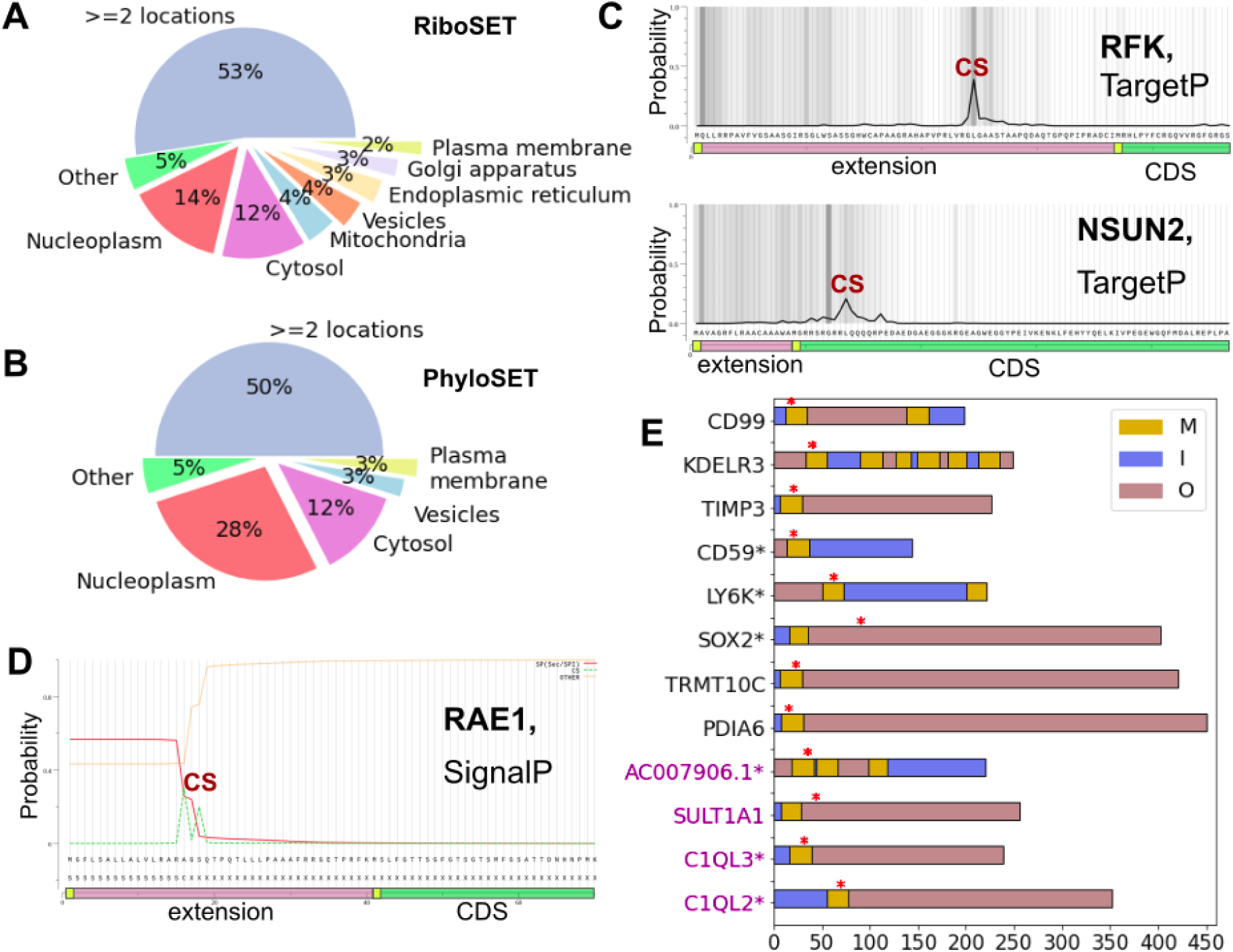
Localisation of PhyloSET and RiboSET genes. Subcellular localisation of RiboSET **(A)** and PhyloSET **(B)** genes according to The Protein Human Atlas. **C.** Probability of mitochondrial presequence calculated by TargetP 2.0 for RFK and NSUN2. Pink bar shows extension, green bar shows 5’ part of CDS, CS - cleavage site. **D.** Probability of signal peptide predicted by SignalP 5.0, for RAE1. Pink bar shows extension, green bar shows 5’ part of CDS, CS - cleavage site. Red line is Signal Peptide, green dotted line is CS (cleavage site), yellow is ‘other’. **E.** Domain organisation of proteforms from RiboSET (black labels) and PhyloSET (purple labels) with TM regions found by TMHMM. Genes supported by Phobius are marked with asterisks. Yellow region (M) is TM helix, blue (I) is inner, pink (O) is outer cell compartment. Red asterisks represent the start of CDS in a proteoform.

One would expect that proteoforms with different alternative N-termini (PANTs) may possess different functional properties. For instance, longer proteoforms may contain a signal of subcellular localization in their extended part for alternative compartmentalisation (Bugler et al. 1991; Tee and Jaffe 2001; Yang et al. 1998). Functional gene enrichment analysis was performed using the Gene Ontology resource (Gene Ontology Consortium 2021; Ashburner et al. 2000) using broader simplified GO terms (Rhee et al. 2008). Significant enrichment for genes from RiboSET was shown in 9 terms in ‘*cellular component*’ and 4 terms in ‘*molecular function*’. We also observed overlaps between terms, e.g. 58 genes associated both with term ’*nucleoplasm*’ and ‘cytoplasm’, 29 common genes between ‘*cytoplasm*’ and ‘*membrane*’ and 16 genes between ‘*nucleoplasm*’ and ‘*membrane*’ terms (Supplementary table 5). No enrichment was shown for PhyloSET genes.

In eukaryotes, N-terminal targeting signals include mitochondrial targeting signal and the signal sequence for the secretory pathway (signal peptides) (Imai and Nakai 2020). Membrane proteins may also contain a signal peptide, but most often the N-terminal transmembrane (TM) region functions as the signal sequence (Johnson and van Waes 1999). First, we extracted the main subcellular location of proteins based on immunofluorescently stained cells from The Human Protein Atlas (HPA) ([CSL STYLE ERROR: reference with no printed form.]; Thul et al. 2017) and for 318 genes from RiboSET Location in HPA was known. The majority of genes are associated with only one compartment (predominantly *Nucleoplasm* -45 genes or *Cytosol -* 39 genes), and for 167 genes two or more localisations are known (Fig.6A, Supplemental table 6). As for PhyloSET, localisation for 40 genes were found in HPA (mostly *Nucleoplasm -* 11 genes and *Cytosol* - 5 genes, 20 genes with 2 or more localisations, Fig.6B, Supplemental table 6).

Next, we utilised algorithms for prediction of localisation signals in the extended proteoforms from RiboSET. SignalP 5.0 is a deep neural network approach that detects the presence of signal peptides and the location of their cleavage sites in proteins (Almagro Armenteros et al. 2019b). We utilised the web-server interface for extended proteins from RiboSET: 29 extended proteoforms have shown the presence of signal peptide and 22 of them have known localisation in HPA. The largest part of them (16) have annotated localisation either in *Golgi apparatus*, *plasma membrane*, *Endoplasmic reticulum* or *vesicles* supporting predictions of SignalP (Supplemental table 6).

Among genes with predicted signal peptides, there was only one gene with detected signal peptide residing entirely within N-terminal extension - *RAE1* (its cleavage site position is situated before the CDS start, location from HPA is ‘*Nucleoli fibrillar centre;Nucleoplasm*’, Fig.6D). Given that signal peptide length varies from 16 aa to 30 aa (von Heijne 1985), for 5 more genes (*STMN1:* ‘Cytosol’*, DPH5:* ‘Golgi apparatus;Nucleoli’*, ADAM15:* unknown*, ZNF622:* ‘Cytosol;Golgi apparatus;Nucleoli;Nucleoplasm’*, SUPT4H1:* ‘Nucleoplasm’) signal peptide spans over CDS and extension. No genes in PhyloSET with signal peptide within theoretical extensions were found (Supplemental table 6).

We also explored the possibility that extended parts of proteoforms can target them to mitochondria. Mitochondrial location was assigned to 20 genes from RiboSET and 2 genes from PhyloSET according to HPA. The length of mitochondrial presequences is 20–60 amino acid residues and they are usually enriched in arginine, leucine, and serine (Calvo et al. 2017; Neupert 1997). We applied TargetP2.0 which is a neural network approach for prediction of various N-terminal presequences including mitochondrial transit peptide (mTP) and found 16 genes in RiboSET with mitochondrial signal (Almagro Armenteros et al. 2019a). Mitochondria is annotated for only 2 genes (*TRMT10C*, *MRPS7*) out of them according to HPA. For 5 genes (*GNAI1: ‘Centrosome;Nucleoli;Nucleoplasm’, RFK: ‘Golgi apparatus’, CDK2: ‘Nucleoplasm;Centrosome;Cytosol’, HSPH1:* ‘*Cytosol;Nucleoplasm’, GNA13: ‘Cytosol*’, Fig.6C) the cleavage site of mitochondrial transfer peptide was located within the extension part. For 6 more genes (*NSUN2: ‘Nucleoplasm’, GCLM: ‘Cytosol;Nucleoplasm;Plasma membrane’, TNKS2: ‘Microtubules’, ACTL6A: ‘Nucleoplasm;Cytosol’, ADRM1: ‘Cytosol;Nucleoplasm;Plasma membrane’, COX7C:* unknown, Fig.6C) the whole mitochondrial signal is likely to span over both extension and CDS according to the minimum length of signal preceding the cleavage site. No mitochondrial presequences were found by TargetP for PhyloSET genes.

Next we explored the existence of N-terminal transmembrane helices within predicted extensions using the transmembrane domain prediction ability of TMHMM (Möller et al. 2001) and Phobius (Käll et al. 2004). We supplemented results from TMHMM with the result of Phobius, because of the issue with high similarity between the hydrophobic regions of a transmembrane helix and that of a signal peptide, leading to cross-reaction between the two types of predictions. The prediction made by *TMHMM* contained 83 genes from RiboSET with TM regions (Supplemental table 6), among which the first TM helix is located at least partially within the extension part in 8 genes, Fig.6E, black labels). According to HPA, only for *LY6K* the location is ‘*Plasma membrane*’ and ‘*Nucleoplasm*’, while *CD99* is assigned with ‘*Golgi apparatus*’, *KDELR3* is in ‘*Endoplasmic reticulum*’, *PDIA6* can be found in ‘*Cytosol*’, ‘*Endoplasmic reticulum*’, *TRMT10C* has ‘*Mitochondria*’ and ‘*Nucleoplasm*’ location and *SOX2* resides in ‘*Nucleoplasm*’. Among PhyloSET genes, 12 were predicted by TMHMM to have a TM in their extended proteoforms. In 4 genes the first TM helix resides at least partly in extension (Fig.6E, purple labels). HPA has no assigned locations for these genes. We supplemented our findings of TM regions with Phobius and found 88 genes containing TM helices among RiboSET genes (75 overlapping genes with TMHMM) and 13 among PhyloSET genes (11 overlapping genes with TMHMM). The first helix starts within extension in 5 genes in RiboSET and 3 genes in PhyloSET (Fig. S10).

### Exclusive non-AUG initiation

One intriguing aspect of non-AUG initiation is that it can be exclusive, which means that unlike in case of PANTs non-AUG initiated proteoform is the only one synthesised from a transcript. This suggests that non-AUG initiation function might be different from production of alternative proteoforms. There are very few known examples of sole non-AUG initiation e.g. *EIF4G2*, *TRPV6*, *TEAD1*, *STIM2* and the reason why non-AUG is preferred evolutionary over AUG has not been elucidated yet. Here we reported several examples of most likely exclusive non-AUG initiation according to their Ribo-seq profiles (Fig.7, Fig.S11 in Supplemental text 1, Supplemental table 6). Most of our examples are in RiboSET exclusively and only CCDC8 is present in both RiboSET and PhyloSET. We also observed that there might be multiple non-AUG translation initiation sites with varying efficiency. For instance, the second potential initiation peak located downstream of predicted non-AUG start in *SLC25A32* (ENST00000297578.8) can be explained by overlap of 5’ leader with another transcript of gene *DCAF13*, located on a different strand (Fig.S12 in Supplemental text 1). However, we cannot exclude the possibility of multiple non-AUG initiations, e.g for the second high peak located upstream of predicted non-AUG start in *THOP1* which may be attributed to GUG codon (Fig.S11 in Supplemental text 1).

**Fig. 7.**
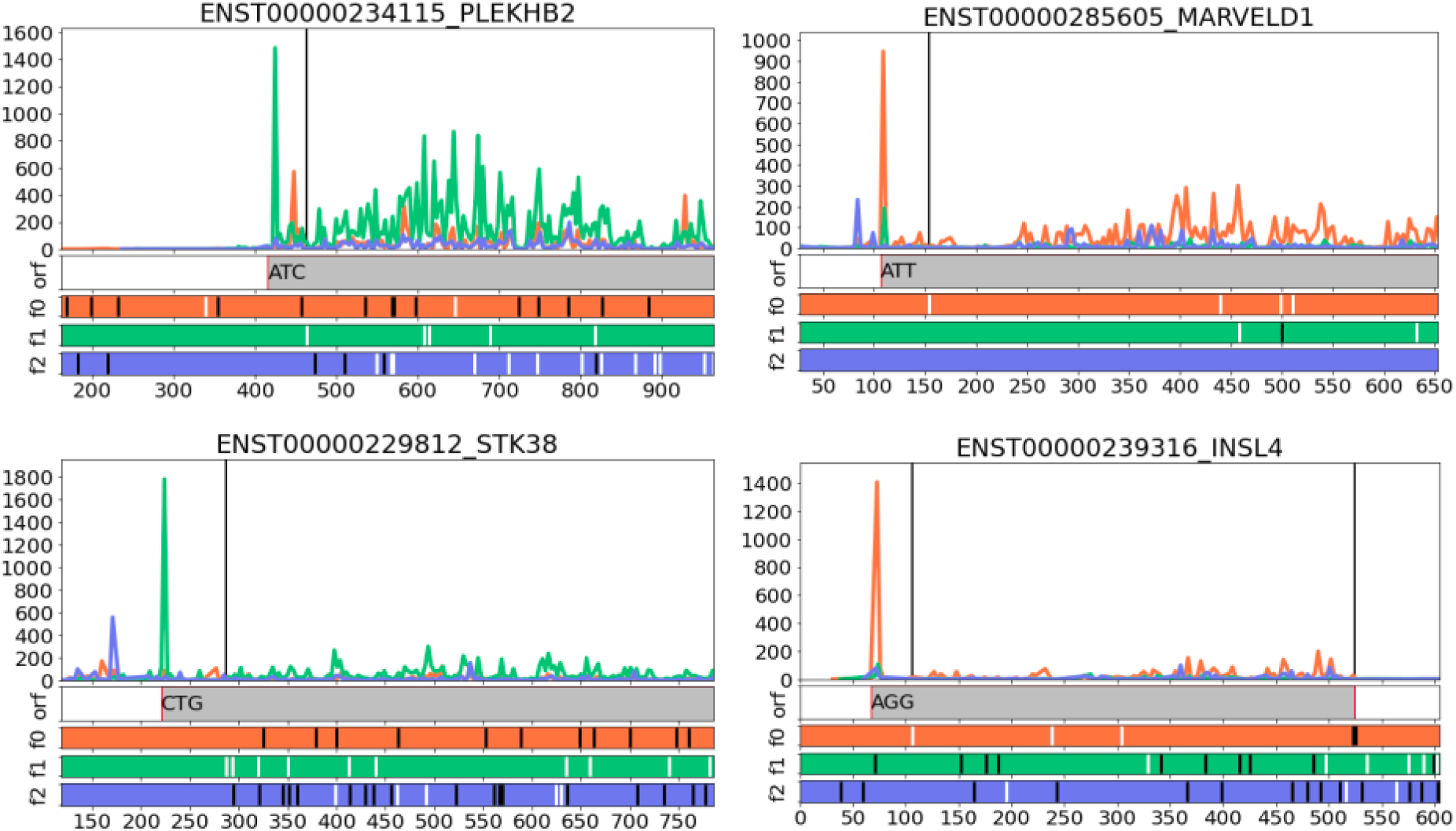
Subcodon Ribo-seq profiles with the densities of ribosome footprints differentially coloured based on the supported reading frame for examples of predicted genes with exclusive non-AUG initiation (*STK38*, *MARVELD1*, *PLEKHB2*, *INSL4*). The colours are matched to the reading frames in ORF plot at the bottom where AUG codons are depicted as white and stop codons as black dashes. Black vertical lines indicate the starts of the annotated CDS. Grey bars correspond to extended CDS initiated at the proposed non-AUG starts. The genomic intervals including primary extension and at least 150 first codons of CDS are shown.

### Reannotation of non_AUG proteoforms

One of the goals of this study is updating human gene annotation with newly identified non-AUG proteoforms as well as reannotation of incomplete or incorrect transcript isoforms discovered along the way. GENCODE also maintains annotation of mouse genes, so human orthologs in mouse have also been incorporated. Of note, annotation requires not only the information about extension being present for that gene, but also an exact position of translation initiation which is challenging to infer precisely due to multiple reasons.

Therefore, the initial phase of annotation includes several immediate cases. Eleven genes have maintained canonical AUG start is their models in human, while additional models with non-AUG start have been introduced: *SFPQ* (CUG), *VANGL2* (AUA), *CCDC8* (CUG), *PELI2* (CUG), *CYTH2* (CUG), *FXR2* (GUG), *H1F* or *H1-10* (CUG), *RPTOR* (CUG), *USP19* (AUA), *SLC6A1* (CUG), *NPLOC4* (GUG). Similarly, non-AUG extensions will be annotated in the next GENCODE release for mice orthologs (*Sfpq*, *Vangl2*, *Ccdc8*, *Pel*, *Cyth2*, *Fxr2*, *H1fx* or *H1f10*, *Rptor*, *Usp19*, *Slc6a1*, *Nploc4*). One more gene (*XRRA1*) which was not included in RiboSET due to its rank (rank 711, below 500 threshold) has also gotten new transcript models both for mouse and human (GUG). Additionally, AUG-extended proteoform in human (*PTPRJ*) has been introduced.

## Discussion

In this study we developed two approaches for detection of non-AUG initiated proteoforms in the human genome. Comparative genomics approach is based on utilisation of the PhyloCSF score which is able to capture signatures of protein-coding evolution in the upstream region of CDS. It resulted in 60 candidate genes (PhyloSET). The second approach employs Ribo-seq data to predict translated extended proteoforms. The analysis of 500 top ranked extensions based on Ribo-seq support led to 392 gene candidates after additional filtering (RiboSET) with only 8 common genes between RiboSET and PhyloSET.

The large overlap between the sets is not expected. For example, some genes from PhyloSET may not be expressed in the cells for which Ribo-seq data are available. Indeed the lower CDS coverage, the lower the extension ranking based on Ribo-seq data, see Fig. 4C. Interestingly, however, we saw several genes with no rank assigned (no Ribo-seq evidence) having significantly high CDS coverage. Given strong evolutionary support for these extensions there could be two potential explanations for this. The extended proteoforms are required in extremely small quantities or they are being synthesised only in specific cells or under specific conditions for which Ribo-seq data were not available. It would be interesting to explore the latter possibility using a larger sets of Ribo-seq data. The correlation between CDS coverage in PhyloSET genes and their Ribo-seq ranking below the top 500 suggests that lower Ribo-seq rankings still correspond to genuine translation. We further tested this possibility by exploring the predictive value of Ribo-seq ranking in finding genes from PhyloSET. Indeed, even the sets of the lowest ranking genes contain more genes in PhyloSET than what is expected by chance (Fig. 3D). Furthermore, around a quarter of extensions in RiboSET and PhyloSET were confirmed with proteomics data available in Trips-Viz (Fig.S1 in Supplementary text 1). Thus, while we cannot estimate the exact number of N-terminal extensions it is likely that it is much greater than 392.

What is the reason for the lack of phylogenetics support for the most N-terminal extensions detectable with Ribo-seq? First N-termini are often processed (Choo et al. 2009; Kunze and Berger 2015), thus it may not matter where exactly initiation occurs, at the annotated AUG codon or at the upstream non-AUG since the mature protein product is the same. The evolution of non-AUG starts upstream of AUG codons is expected to be neutral in this case and would not be detected with the PhyloCSF approach. Similar situation would be expected when N-terminal extensions do not alter the functional properties of the proteins. Nonetheless, it does not mean that all extensions occurring exclusively in RiboSET are inconsequential. First, they may shape the immune response by providing potential antigens. Second, mutations in the regions corresponding to N-terminal extensions may have different effects from those occurring in non-coding parts of 5’ leaders. For example, an introduction of an in-frame stop codon would result in an uORF that would inhibit translation of CDS. It is also conceivable that in a number of cases the evolutionary selection was not detected with our approach. Some extensions may be very recent, so that there is insufficient number of substitutions in the narrow phylogenetic group in which they exist, some may even be uniquely human. The analysis of the data in aggregated databases of personal genomes, such as gnomAD may eventually reveal evolutionary selection in such cases. Also, we may have failed to detect evolutionary selection in those cases where it acts only at a very short region, below 50 codons upstream that we used in this study.

Of note, positive PhyloCSF score upstream of annotated starts may have a different reason than non-AUG extension. There is a possibility that positive PhyloCSF signal might be explained by the remnant of CDS which was truncated in humans by nonsense mutation. For instance, in *LRP5L*, nonsense mutation happened in the ancestor of human and chimpanzee thus possibly disrupting a CDS present in other mammals (Supplemental Fig 14). Therefore the next in-frame AUG which was internal became a new start in humans. Nevertheless, it does not exclude the possibility that human-specific non-AUG extensions can exist and be translated under certain conditions. There is also a need for acknowledgement of a limitation of genome annotations. For example, 5’ends of transcript may be incomplete. For *HES3* gene, in GENCODE v38 there is an upstream AUG which has a clearly Ribo-seq stalling signal according to GWIPs-viz and it is well-conserved in mammals. However, in GENCODE v25 (that we used) 5’ends of *HES3* transcripts are shorter and do not contain such AUG thus making it impossible to detect such AUG.

Interestingly, one of the possible functions of non-AUG extension could be alternative compartmentalisation. GO analysis has shown that there are plenty of common genes between subcellular localisation terms. It can be interpreted as there are multiple subcellular localisations for gene products which might be facilitated by extensions. Also, data from HPA where 167 genes can be found in multiple compartments also leads to an idea that such distinct locations may correspond to proteoforms with alternative N-termini. This led us to applying tools for predicting signal peptides, mitochondrial presequences and transmembrane domains which revealed that there are genes with localisation signals residing within their non-AUG extensions.

As a result of these study we reannotated 11 human genes and also provided information on likely functions of some of these extensions in differential compartmentalisation of short and long proteoforms.

## Methods

### A pipeline for detection of non-AUG proteoforms using evolutionary signatures

We obtained 94359 human protein-coding mRNA sequences from GENCODE v25 (GRCh38.p7), this particular version was chosen because of the available processed Ribo-seq and proteomics data available in the Trips-Viz browser (Kiniry et al. 2019). For each transcript starting from an annotated AUG codon we moved along the transcript in the 5’ direction by 3 nucleotides until an in-frame STOP codon was reached. We termed this sequence primary (or theoretical) extension. For assessing PhyloCSF score (Lin et al. 2011) using 120 mammals alignment in .maf format (Hecker and Hiller 2020) we took 50 codons upstream of the annotated AUG (or less if length of primary extension is shorter, genes where theoretical extension is less 20 codons are discarded). MafExtract from CESAR2.0 (Sharma et al. 2017) coupled with PHAST (Hubisz et al. 2011) and custom Python scripts were used to extract multiple sequence alignments for 50 codons of N-termini. Transcripts with a positive PhyloCSF score for 50-codons-long or shorter N termini (3058 genes and 5417 transcripts) were selected as candidates for further analysis.

We excluded transcripts for which theoretical N-terminal extensions have any overlaps with coding exons in the same strand from GENCODE v25 release (GRCh38.p7), GENCODE v35 release (GRCh38.p13), RefSeq (July 1, 2020; GRCh38.p13, 109.20200815). Briefly for GENCODE annotations we extracted transcriptomic coordinates of CDS from protein-coding fasta files (gencode.v25.pc_transcripts.fa, gencode.v35.pc_transcripts.fa) and transformed to genomic coordinates using pmapFromTranscripts from the GenomicFeatures package in R (v3.6.1). For RefSeq annotations coding exons were extracted from GRCh38_latest_genomic.gff. Intersections were identified using bedtools intersect (Quinlan and Hall 2010) (v2.29.2).

### Detection of translated regions in Trips-Viz

We used the corresponding Trips-Viz functionality using Ribo-seq data from 13 studies GSE62247 (Werner et al. 2015), GSE114794 (Gameiro and Struhl 2018), GSE79664 (Park et al. 2016), GSE51584 (Guo et al. 2014), GSE94460 (Zhang et al. 2017), GSE73136 (Calviello et al. 2016), GSE87328 (Fijalkowska et al. 2017), GSE64962 (Xu et al. 2016), GSE65885 (Ji et al. 2016), GSE56887 (Wolfe et al. 2014), GSE70211 and GSE79392 (Iwasaki et al. 2016), GSE77401 (Goodarzi et al. 2016), GSE58207 (Crappé et al. 2015) and 4 studies of proteomics data PXD004452 (Bekker-Jensen et al. 2017), PXD002395 (Geiger et al. 2012), PXD002082 (Zhang et al. 2014), PXD002815 (Hein et al. 2015).

In the publicly available version of Trips-viz only AUG, CUG and GUG-initiated extensions are available to retrieve, so additional non-cognate starts including UUG, AUA, AUU, AUC, ACG, AGG, AAG were supplied into the in-house Trips-viz version. The top-500 ranked extensions were used to construct RiboSET. The Trips-Viz ranking procedure of extended proteoforms includes two steps: (1) calculation of three parameters (Highest Reading Frame, StartRiseUp, Non-Zero Coverage) for each selected non-AUG extension; (2) final ranking is produced by summing the ranks of the individual features. More detailed description can be found in Supplementary text 2.

### Characterisation of predicted non-AUG starts in RiboSET

TIS scores were extracted from (Diaz de Arce et al. 2018). For RNA secondary structure analysis 400 genes initiating with AUG codon with the highest TIS-scores and the lowest TIS-scores were selected for comparison from RiboSET. Free Gibbs energy (dG) was calculated using RNAfold 2.4.17 (Lorenz et al. 2011) within a sliding window of 22nt at 1nt step in the region starting from 10nt upstream and up to 100nt downstream of the start codon (averaged by genes with 95% confidence interval using Python 3.7.3 package statsmodels v 0.10.1). Multiple sequence alignment was retrieved by MAFFT v7.310 (Katoh and Standley 2013) and plotted by ggmsa_1.0.0, R v3.6.1. Sequence logo was built by ggseqlogo_0.1, R v3.6.1. Data and all other plots were analysed by pandas 1.2.1 and drawn using matplotlib 3.2.1, Python 3.7.3. GO enrichment analysis on RiboSET and PhyloSET genes was performed using GOATOOLS, v1.1.6 (Klopfenstein et al. 2018) with GO-basic version of database ([CSL STYLE ERROR: reference with no printed form.]) and GENCODE v25 protein coding genes as reference set, significant terms were selected based on adjusted p-value < 0.05 (after using fdr multiple testing correction).

As a source of experimentally supported protein localisation The Human Protein Atlas was used. In order to predict signal peptides we used SignalP 5.0 (Almagro Armenteros et al. 2019b), for identification of mitochondrial presequences we applied TargetP 2.0 (Almagro Armenteros et al. 2019a) and for extraction of TM domains Phobius (Käll et al. 2007) and TMHMM (Möller et al. 2001) were employed.

### Data access

All data generated during this study are included in this published article and its supplementary information files: Supplemental file 1 (Supplemental_text_1.docx), Supplemental file 2 (Supplementary_text_2.docx), Supplemental table 1 (Supplemental_table_S1.xslx), Supplemental table 2 (Supplemental_table_S2.xslx), Supplemental table 3 (Supplemental_table_S3.xslx), Supplemental table 4 (Supplemental_table_S4.xslx), Supplemental table 5 (Supplemental_table_S5.xslx), Supplemental table 6 (Supplemental_table_S6.xslx), Supplemental table 7 (Supplemental_table_S7.xslx), Supplemental table 8 (Supplemental_table_S.xslx). The datasets analysed during the current study are available in GEO under accession numbers: GSE62247, GSE114794, GSE79664, GSE51584, GSE94460, GSE73136, GSE87328, GSE64962, GSE65885, GSE56887, GSE70211, GSE79392, GSE77401, GSE58207, GSE111891, GSE83332 and in the ProteomeXchange: PXD004452, PXD002395, PXD002082, PXD002815. Custom code can be found in https://github.com/triasteran/nonAUG_manuscript/tree/main/jupyter_notebooks.

## Acknowledgements

We are grateful to Irwin Jungreis and Manolis Kellis at CSAIL (MIT) for supplying PhyloCSF parameters for 120-way mammalian genomic alignment and making CodAlignView available to us. This work was supported by SFI-HRB-Wellcome Trust Biomedical Research Partnership (Investigator Award in Science) [210692] to P.V.B; Science Foundation Ireland Centre for Research Training in Genomics Data Science (18/CRT/6214 to A.D.F) and Russian Science Foundation (20-14-00121) to D.E.A. S.J.K. wishes to acknowledge support from the Irish Research Council. JMM is supported by the National Human Genome Research Institute of the National Institutes of Health under award number 2U41HG007234 and the European Molecular Biology Laboratory. For the purpose of open access, the author has applied a CC BY public copyright licence to any Author Accepted Manuscript version arising from this submission. The content is solely the responsibility of the authors and does not necessarily represent the official views of the National Institutes of Health. Ensembl is a registered trademark of EMBL.

P.V.B conceived the work, secured funding and supervised the study. A.D.F performed most of the computational analyses, prepared the first draft of the manuscript and all data figures. S.J.K implemented the ranking algorithm for identification of translated non-AUGs in Trips-Viz. J.M.M. evaluated the suitability of N-terminal extension for relevant GENCODE reannotations. All authors participated in interpretation of the data and editing the final version of the manuscript.

## References

Almagro Armenteros JJ, Salvatore M, Emanuelsson O, Winther O, von Heijne G, Elofsson A, Nielsen H. 2019a. Detecting sequence signals in targeting peptides using deep learning. Life Sci Alliance 2. http://dx.doi.org/10.26508/lsa.201900429.

Almagro Armenteros JJ, Tsirigos KD, Sønderby CK, Petersen TN, Winther O, Brunak S, von Heijne G, Nielsen H. 2019b. SignalP 5.0 improves signal peptide predictions using deep neural networks. Nat Biotechnol 37: 420–423.

Anderson CW, Buzash-Pollert E. 1985. Can ACG serve as an initiation codon for protein synthesis in eucaryotic cells? Mol Cell Biol 5: 3621–3624.

Andreev DE, O’Connor PBF, Fahey C, Kenny EM, Terenin IM, Dmitriev SE, Cormican P, Morris DW, Shatsky IN, Baranov PV. 2015a. Translation of 5’ leaders is pervasive in genes resistant to eIF2 repression. Elife 4: e03971.

Andreev DE, O’Connor PBF, Zhdanov AV, Dmitriev RI, Shatsky IN, Papkovsky DB, Baranov PV. 2015b. Oxygen and glucose deprivation induces widespread alterations in mRNA translation within 20 minutes. Genome Biol 16: 90.

Arnaud E, Touriol C, Boutonnet C, Gensac MC, Vagner S, Prats H, Prats AC. 1999. A new 34-kilodalton isoform of human fibroblast growth factor 2 is cap dependently synthesized by using a non-AUG start codon and behaves as a survival factor. Mol Cell Biol 19: 505–514.

Ashburner M, Ball CA, Blake JA, Botstein D, Butler H, Cherry JM, Davis AP, Dolinski K, Dwight SS, Eppig JT, et al. 2000. Gene ontology: tool for the unification of biology. The Gene Ontology Consortium. Nat Genet 25: 25–29.

Atkins JF, Loughran G, Bhatt PR, Firth AE, Baranov PV. 2016. Ribosomal frameshifting and transcriptional slippage: From genetic steganography and cryptography to adventitious use. Nucleic Acids Res 44: 7007–7078.

Baranov PV, Atkins JF, Yordanova MM. 2015. Augmented genetic decoding: global, local and temporal alterations of decoding processes and codon meaning. Nat Rev Genet 16: 517–529.

Baranov PV, Gesteland RF, Atkins JF. 2004. P-site tRNA is a crucial initiator of ribosomal frameshifting. RNA 10: 221–230.

Bekker-Jensen DB, Kelstrup CD, Batth TS, Larsen SC, Haldrup C, Bramsen JB, Sørensen KD, Høyer S, Ørntoft TF, Andersen CL, et al. 2017. An Optimized Shotgun Strategy for the Rapid Generation of Comprehensive Human Proteomes. Cell Syst 4: 587–599.e4.

Bugler B, Amalric F, Prats H. 1991. Alternative initiation of translation determines cytoplasmic or nuclear localization of basic fibroblast growth factor. Mol Cell Biol 11: 573–577.

Calviello L, Mukherjee N, Wyler E, Zauber H, Hirsekorn A, Selbach M, Landthaler M, Obermayer B, Ohler U. 2016. Detecting actively translated open reading frames in ribosome profiling data. Nat Methods 13: 165–170.

Calvo SE, Julien O, Clauser KR, Shen H, Kamer KJ, Wells JA, Mootha VK. 2017. Comparative Analysis of Mitochondrial N-Termini from Mouse, Human, and Yeast. Mol Cell Proteomics 16: 512–523.

Choo KH, Tan TW, Ranganathan S. 2009. A comprehensive assessment of N-terminal signal peptides prediction methods. BMC Bioinformatics 10 **Suppl 15**: S2.

Crappé J, Ndah E, Koch A, Steyaert S, Gawron D, De Keulenaer S, De Meester E, De Meyer T, Van Criekinge W, Van Damme P, et al. 2015. PROTEOFORMER: deep proteome coverage through ribosome profiling and MS integration. Nucleic Acids Res 43: e29.

Diaz de Arce AJ, Noderer WL, Wang CL. 2018. Complete motif analysis of sequence requirements for translation initiation at non-AUG start codons. Nucleic Acids Res 46: 985–994.

Fijalkowska D, Verbruggen S, Ndah E, Jonckheere V, Menschaert G, Van Damme P. 2017. eIF1 modulates the recognition of suboptimal translation initiation sites and steers gene expression via uORFs. Nucleic Acids Res 45: 7997–8013.

Gameiro PA, Struhl K. 2018. Nutrient Deprivation Elicits a Transcriptional and Translational Inflammatory Response Coupled to Decreased Protein Synthesis. Cell Rep 24: 1415– 1424.

Gallant J, Lindsley D, Masucci J. The Ribosome: Structure, Function, Antibiotics, and Cellular Interactions, chapter 31. The Unbearable Lightness of Peptidyl-tRNA.

Geiger T, Wehner A, Schaab C, Cox J, Mann M. 2012. Comparative proteomic analysis of eleven common cell lines reveals ubiquitous but varying expression of most proteins. Mol Cell Proteomics 11: M111.014050.

Gene Ontology Consortium. 2021. The Gene Ontology resource: enriching a GOld mine. Nucleic Acids Res 49: D325–D334.

Goodarzi H, Nguyen HCB, Zhang S, Dill BD, Molina H, Tavazoie SF. 2016. Modulated Expression of Specific tRNAs Drives Gene Expression and Cancer Progression. Cell 165: 1416–1427.

Gruber AR, Lorenz R, Bernhart SH, Neuböck R, Hofacker IL. 2008. The Vienna RNA websuite. Nucleic Acids Res 36: W70–4.

Gunišová S, Hronová V, Mohammad MP, Hinnebusch AG, Valášek LS. 2018. Please do not recycle! Translation reinitiation in microbes and higher eukaryotes. FEMS Microbiol Rev 42: 165–192.

Guo JU, Agarwal V, Guo H, Bartel DP. 2014. Expanded identification and characterization of mammalian circular RNAs. Genome Biol 15: 409.

Hann SR, Eisenman RN. 1984. Proteins encoded by the human c-myc oncogene: differential expression in neoplastic cells. Mol Cell Biol 4: 2486–2497.

Hann SR, King MW, Bentley DL, Anderson CW, Eisenman RN. 1988. A non-AUG translational initiation in c-myc exon 1 generates an N-terminally distinct protein whose synthesis is disrupted in Burkitt’s lymphomas. Cell 52: 185–195.

Hecker N, Hiller M. 2020. A genome alignment of 120 mammals highlights ultraconserved element variability and placenta-associated enhancers. Gigascience 9. http://dx.doi.org/10.1093/gigascience/giz159.

Hein MY, Hubner NC, Poser I, Cox J, Nagaraj N, Toyoda Y, Gak IA, Weisswange I, Mansfeld J, Buchholz F, et al. 2015. A human interactome in three quantitative dimensions organized by stoichiometries and abundances. Cell 163: 712–723.

Hinnebusch AG. 2014. The scanning mechanism of eukaryotic translation initiation. Annu Rev Biochem 83: 779–812.

Hopkins BD, Fine B, Steinbach N, Dendy M, Rapp Z, Shaw J, Pappas K, Yu JS, Hodakoski C, Mense S, et al. 2013. A secreted PTEN phosphatase that enters cells to alter signaling and survival. Science 341: 399–402.

Hubisz MJ, Pollard KS, Siepel A. 2011. PHAST and RPHAST: phylogenetic analysis with space/time models. Brief Bioinform 12: 41–51.

Imai K, Nakai K. 2020. Tools for the Recognition of Sorting Signals and the Prediction of Subcellular Localization of Proteins From Their Amino Acid Sequences. Front Genet 11: 607812.

Imataka H, Olsen HS, Sonenberg N. 1997. A new translational regulator with homology to eukaryotic translation initiation factor 4G. EMBO J 16: 817–825.

Ingolia NT, Brar GA, Rouskin S, McGeachy AM, Weissman JS. 2012. The ribosome profiling strategy for monitoring translation in vivo by deep sequencing of ribosome-protected mRNA fragments. Nat Protoc 7: 1534–1550.

Ivanov IP, Firth AE, Michel AM, Atkins JF, Baranov PV. 2011. Identification of evolutionarily conserved non-AUG-initiated N-terminal extensions in human coding sequences. Nucleic Acids Res 39: 4220–4234.

Ivanov IP, Gaikwad S, Hinnebusch AG, Dever TE. 2020. Conserved +1 translational frameshifting in the S. cerevisiae gene encoding YPL034W. bioRxiv 2020.04.29.069534. https://www.biorxiv.org/content/10.1101/2020.04.29.069534v1.abstract (Accessed January 19, 2022).

Ivanov IP, Saba JA, Fan C-M, Wang J, Firth AE, Cao C, Green R, Dever TE. 2022. Evolutionarily conserved inhibitory uORFs sensitize mRNA translation to start codon selection stringency. Proc Natl Acad Sci U S A 119. http://dx.doi.org/10.1073/pnas.2117226119.

Iwasaki S, Floor SN, Ingolia NT. 2016. Rocaglates convert DEAD-box protein eIF4A into a sequence-selective translational repressor. Nature 534: 558–561.

Ji Z, Song R, Huang H, Regev A, Struhl K. 2016. Transcriptome-scale RNase-footprinting of RNA-protein complexes. Nat Biotechnol 34: 410–413.

Johnson AE, van Waes MA. 1999. The translocon: a dynamic gateway at the ER membrane. Annu Rev Cell Dev Biol 15: 799–842.

Käll L, Krogh A, Sonnhammer ELL. 2004. A combined transmembrane topology and signal peptide prediction method. J Mol Biol 338: 1027–1036.

Käll L, Krogh A, Sonnhammer ELL. 2007. Advantages of combined transmembrane topology and signal peptide prediction--the Phobius web server. Nucleic Acids Res 35: W429– 32.

Karagyozov L, Grozdanov PN, Böhmer F-D. 2020. The translation attenuating arginine-rich sequence in the extended signal peptide of the protein-tyrosine phosphatase PTPRJ/DEP1 is conserved in mammals. PLoS One 15: e0240498.

Katoh K, Standley DM. 2013. MAFFT multiple sequence alignment software version 7: improvements in performance and usability. Mol Biol Evol 30: 772–780.

Kearse MG, Green KM, Krans A, Rodriguez CM, Linsalata AE, Goldstrohm AC, Todd PK. 2016. CGG Repeat-Associated Non-AUG Translation Utilizes a Cap-Dependent Scanning Mechanism of Initiation to Produce Toxic Proteins. Mol Cell 62: 314–322.

Kearse MG, Wilusz JE. 2017. Non-AUG translation: a new start for protein synthesis in eukaryotes. Genes Dev 31: 1717–1731.

Khan YA, Jungreis I, Wright JC, Mudge JM, Choudhary JS, Firth AE, Kellis M. 2020. Evidence for a novel overlapping coding sequence in POLG initiated at a CUG start codon. BMC Genet 21: 25.

Kiniry SJ, Judge CE, Michel AM, Baranov PV. 2021. Trips-Viz: an environment for the analysis of public and user-generated ribosome profiling data. Nucleic Acids Res 49: W662–W670.

Kiniry SJ, O’Connor PBF, Michel AM, Baranov PV. 2019. Trips-Viz: a transcriptome browser for exploring Ribo-Seq data. Nucleic Acids Res 47: D847–D852.

Klopfenstein DV, Zhang L, Pedersen BS, Ramírez F, Warwick Vesztrocy A, Naldi A, Mungall CJ, Yunes JM, Botvinnik O, Weigel M, et al. 2018. GOATOOLS: A Python library for Gene Ontology analyses. Sci Rep 8: 10872.

Kochetov AV, Palyanov A, Titov II, Grigorovich D, Sarai A, Kolchanov NA. 2007. AUG_hairpin: prediction of a downstream secondary structure influencing the recognition of a translation start site. BMC Bioinformatics 8: 318.

Kozak M. 1989. Context effects and inefficient initiation at non-AUG codons in eucaryotic cell-free translation systems. Mol Cell Biol 9: 5073–5080.

Kozak M. 1990. Downstream secondary structure facilitates recognition of initiator codons by eukaryotic ribosomes. Proc Natl Acad Sci U S A 87: 8301–8305.

Kozak M. 1980. Evaluation of the “scanning model” for initiation of protein synthesis in eucaryotes. Cell 22: 7–8.

Kunze M, Berger J. 2015. The similarity between N-terminal targeting signals for protein import into different organelles and its evolutionary relevance. Front Physiol 6: 259.

Liang H, Chen X, Yin Q, Ruan D, Zhao X, Zhang C, McNutt MA, Yin Y. 2017. PTENβ is an alternatively translated isoform of PTEN that regulates rDNA transcription. Nat Commun 8: 14771.

Lin MF, Jungreis I, Kellis M. 2011. PhyloCSF: a comparative genomics method to distinguish protein coding and non-coding regions. Bioinformatics 27: i275–82.

Lorenz R, Bernhart SH, Höner Zu Siederdissen C, Tafer H, Flamm C, Stadler PF, Hofacker IL. 2011. ViennaRNA Package 2.0. Algorithms Mol Biol 6: 26.

Loughran G, Howard MT, Firth AE, Atkins JF. 2017. Avoidance of reporter assay distortions from fused dual reporters. RNA 23: 1285–1289.

Loughran G, Zhdanov AV, Mikhaylova MS, Rozov FN, Datskevich PN, Kovalchuk SI, Serebryakova MV, Kiniry SJ, Michel AM, O’Connor PBF, et al. 2020. Unusually efficient CUG initiation of an overlapping reading frame in mRNA yields novel protein POLGARF. Proc Natl Acad Sci U S A 117: 24936–24946.

Michel AM, Fox G, M Kiran A, De Bo C, O’Connor PBF, Heaphy SM, Mullan JPA, Donohue CA, Higgins DG, Baranov PV. 2014. GWIPS-viz: development of a ribo-seq genome browser. Nucleic Acids Res 42: D859–64.

Möller S, Croning MD, Apweiler R. 2001. Evaluation of methods for the prediction of membrane spanning regions. Bioinformatics 17: 646–653.

Mueller PP, Hinnebusch AG. 1986. Multiple upstream AUG codons mediate translational control of GCN4. Cell 45: 201–207.

Neupert W. 1997. Protein import into mitochondria. Annu Rev Biochem 66: 863–917.

Ogle JM, Brodersen DE, Clemons WM Jr, Tarry MJ, Carter AP, Ramakrishnan V. 2001. Recognition of cognate transfer RNA by the 30S ribosomal subunit. Science 292: 897–902.

Palanimurugan R, Scheel H, Hofmann K, Dohmen RJ. 2004. Polyamines regulate their synthesis by inducing expression and blocking degradation of ODC antizyme. EMBO J 23: 4857–4867.

Park J-E, Yi H, Kim Y, Chang H, Kim VN. 2016. Regulation of Poly(A) Tail and Translation during the Somatic Cell Cycle. Mol Cell 62: 462–471.

Peabody DS. 1989. Translation initiation at non-AUG triplets in mammalian cells. J Biol Chem 264: 5031–5035.

Potapov AP, Triana-Alonso FJ, Nierhaus KH. 1995. Ribosomal decoding processes at codons in the A or P sites depend differently on 2’-OH groups. J Biol Chem 270: 17680–17684.

Quinlan AR, Hall IM. 2010. BEDTools: a flexible suite of utilities for comparing genomic features. Bioinformatics 26: 841–842.

Ramakrishnan V. 2002. Ribosome structure and the mechanism of translation. Cell 108: 557–572.

Reuter K, Biehl A, Koch L, Helms V. 2016. PreTIS: A Tool to Predict Non-canonical 5’ UTR Translational Initiation Sites in Human and Mouse. PLoS Comput Biol 12: e1005170.

Rhee SY, Wood V, Dolinski K, Draghici S. 2008. Use and misuse of the gene ontology annotations. Nat Rev Genet 9: 509–515.

Sharma V, Schwede P, Hiller M. 2017. CESAR 2.0 substantially improves speed and accuracy of comparative gene annotation. Bioinformatics 33: 3985–3987.

Simonetti A, Marzi S, Myasnikov AG, Fabbretti A, Yusupov M, Gualerzi CO, Klaholz BP. 2008. Structure of the 30S translation initiation complex. Nature 455: 416–420.

Skabkin MA, Skabkina OV, Hellen CUT, Pestova TV. 2013. Reinitiation and other unconventional posttermination events during eukaryotic translation. Mol Cell 51: 249– 264.

Starck SR, Tsai JC, Chen K, Shodiya M, Wang L, Yahiro K, Martins-Green M, Shastri N, Walter P. 2016. Translation from the 5’ untranslated region shapes the integrated stress response. Science 351: aad3867.

Svidritskiy E, Korostelev AA. 2015. Ribosome Structure Reveals Preservation of Active Sites in the Presence of a P-Site Wobble Mismatch. Structure 23: 2155–2161.

Tailor CS, Marin M, Nouri A, Kavanaugh MP, Kabat D. 2001. Truncated forms of the dual function human ASCT2 neutral amino acid transporter/retroviral receptor are translationally initiated at multiple alternative CUG and GUG codons. J Biol Chem 276: 27221–27230.

Takahashi K, Maruyama M, Tokuzawa Y, Murakami M, Oda Y, Yoshikane N, Makabe KW, Ichisaka T, Yamanaka S. 2005. Evolutionarily conserved non-AUG translation initiation in NAT1/p97/DAP5 (EIF4G2). Genomics 85: 360–371.

Tang L, Morris J, Wan J, Moore C, Fujita Y, Gillaspie S, Aube E, Nanda J, Marques M, Jangal M, et al. 2017. Competition between translation initiation factor eIF5 and its mimic protein 5MP determines non-AUG initiation rate genome-wide. Nucleic Acids Res 45: 11941–11953.

Tee MK, Jaffe RB. 2001. A precursor form of vascular endothelial growth factor arises by initiation from an upstream in-frame CUG codon. Biochem J 359: 219–226.

Thul PJ, Åkesson L, Wiking M, Mahdessian D, Geladaki A, Ait Blal H, Alm T, Asplund A, Björk L, Breckels LM, et al. 2017. A subcellular map of the human proteome. Science 356. http://dx.doi.org/10.1126/science.aal3321.

Tzani I, Ivanov IP, Andreev DE, Dmitriev RI, Dean KA, Baranov PV, Atkins JF, Loughran G. 2016. Systematic analysis of the PTEN 5’ leader identifies a major AUU initiated proteoform. Open Biol 6. http://dx.doi.org/10.1098/rsob.150203.

Van Damme P, Gawron D, Van Criekinge W, Menschaert G. 2014. N-terminal proteomics and ribosome profiling provide a comprehensive view of the alternative translation initiation landscape in mice and men. Mol Cell Proteomics 13: 1245–1261.

von Heijne G. 1985. Signal sequences. The limits of variation. J Mol Biol 184: 99–105.

Werner A, Iwasaki S, McGourty CA, Medina-Ruiz S, Teerikorpi N, Fedrigo I, Ingolia NT, Rape M. 2015. Cell-fate determination by ubiquitin-dependent regulation of translation. Nature 525: 523–527.

Williams RT, Manji SS, Parker NJ, Hancock MS, Van Stekelenburg L, Eid JP, Senior PV, Kazenwadel JS, Shandala T, Saint R, et al. 2001. Identification and characterization of the STIM (stromal interaction molecule) gene family: coding for a novel class of transmembrane proteins. Biochem J 357: 673–685.

Wolfe AL, Singh K, Zhong Y, Drewe P, Rajasekhar VK, Sanghvi VR, Mavrakis KJ, Jiang M, Roderick JE, Van der Meulen J, et al. 2014. RNA G-quadruplexes cause eIF4A-dependent oncogene translation in cancer. Nature 513: 65–70.

Xiao JH, Davidson I, Matthes H, Garnier JM, Chambon P. 1991. Cloning, expression, and transcriptional properties of the human enhancer factor TEF-1. Cell 65: 551–568.

Xu B, Gogol M, Gaudenz K, Gerton JL. 2016. Improved transcription and translation with L-leucine stimulation of mTORC1 in Roberts syndrome. BMC Genomics 17: 25.

Yang X, Chernenko G, Hao Y, Ding Z, Pater MM, Pater A, Tang SC. 1998. Human BAG-1/RAP46 protein is generated as four isoforms by alternative translation initiation and overexpressed in cancer cells. Oncogene 17: 981–989.

Yeom J, Ju S, Choi Y, Paek E, Lee C. 2017. Comprehensive analysis of human protein N-termini enables assessment of various protein forms. Sci Rep 7: 6599.

Zhang B, Wang J, Wang X, Zhu J, Liu Q, Shi Z, Chambers MC, Zimmerman LJ, Shaddox KF, Kim S, et al. 2014. Proteogenomic characterization of human colon and rectal cancer. Nature 513: 382–387.

Zhang P, He D, Xu Y, Hou J, Pan B-F, Wang Y, Liu T, Davis CM, Ehli EA, Tan L, et al. 2017. Genome-wide identification and differential analysis of translational initiation. Nat Commun 8: 1749.

Zhang X, Gao X, Coots RA, Conn CS, Liu B, Qian S-B. 2015. Translational control of the cytosolic stress response by mitochondrial ribosomal protein L18. Nat Struct Mol Biol 22: 404–410.

Download ontology. *Gene Ontology Resource*. http://geneontology.org/docs/download-ontology/ (Accessed January 19, 2022a).

The Human Protein Atlas. http://www.proteinatlas.org (Accessed February 18, 2022b).

